# Mutually exclusive locales for N-linked glycans and disorder in glycoproteins

**DOI:** 10.1101/443143

**Authors:** Alvina Singh, Indu Kumari, Dharma Pally, Shyamili Goutham, Sujasha Ghosh, Yusuf Akhter, Ramray Bhat

## Abstract

Several post-translational modifications of proteins lie within regions of disorder, stretches of amino acid residues that exhibit a dynamic tertiary structure and resist crystallization. Such localization has been proposed to expand the binding versatility of the disordered regions, and hence, the repertoire of interacting partners for the proteins. However, investigating a dataset of 500 human N-linked glycoproteins, we observed that the sites of N-linked glycosylations, or N-glycosites, lay predominantly within the regions of predicted order rather than their unstructured counterparts. This mutual exclusivity between disordered stretches and N-glycosites could not be reconciled merely through asymmetry in distribution of asparagines, serines or threonines residues, which comprise the minimum-required signature for conjugation by N-linked glycans, but rather by a contextual enrichment of these residues next to each other within the ordered portions. In fact, N-glycosite neighborhoods and disordered stretches showed distinct sets of enriched residues suggesting their individualized roles in protein phenotype. N-glycosite neighborhood residues also showed higher phylogenetic conservation than disordered stretches within amniote orthologs of glycoproteins. However, a universal search for residue-combinations that are putatively domain-constitutive ranked the disordered regions higher than the N-glycosite neighborhoods. We propose that amino acid residue-combinations bias the permissivity for N-glycoconjugation within ordered regions, so as to balance the tradeoff between the evolution of protein stability, and function, contributed by the N-linked glycans and disordered regions respectively.

## Introduction

One of most common co- and post-translational modifications of proteins is the conjugation of branched glycosylations to their asparagines (known as N-linked glycosylations, to distinguish them from serine-attached O-linked glycosylation) (Varki, 2009). The process of glycosylation takes place in the rough endoplasmic reticulum and continues in the Golgi complex. Therefore, several proteins that end up in the extracellular milieu or within the transmembrane compartment are glycoconjugates (the converse is not necessarily true: O-GlcNAcylation is an important modification seen in the nucleus (West and Hart, 2015)). The establishment of organismal morphologies has been sought to be understood through the interactions of a highly conserved set of proteins known as the developmental genetic toolkit {Jaramillo, 2016 #56}. Most toolkit proteins, which are involved in tissue-scale processes, such as cell-cell and cell-matrix adhesion, diffusion-driven signaling and cell movement, are extracellular- or membrane-bound glycoproteins (Engler et al., 2009; Newman and Bhat, 2009)) Given their crucial developmental role, it is reasonable to hypothesize that there is an evolutionary constraint on the structure and function of these proteins canalizing their ability to perform discrete roles in tissue- and organ-development.

It is now well known that a large number of eukaryotic proteins show flexible tertiary structures, and are known as intrinsically disordered proteins (IDPs) (Romero et al., 2004; Uversky and Dunker, 2008, 2010). Other proteins, while not being entirely disordered can possess variable lengths of disordered residues. The inherently flexible nature of disordered peptide stretches enhances the proteins’ repertoire of interacting or binding partners, and also plays an important role in the catalytic activity of the protein (Fuxreiter, 2012; Mohan et al., 2006). Multiple cytoplasmic and nuclear proteins (proteins involved in signal transduction and transcription factors) are highly disordered although extracellular proteins have been found to also have variable disordered stretches (Cortese et al., 2008; Dunker and Uversky, 2008; Liu et al., 2006; Niklas et al., 2015). Several post translational modifications (PTMs), which modulate their binding and interactive properties, have been reported to be located within disordered regions e.g. phosphorylations of serines and threonines are found within unstructured regions both in eukaryotes (Amoutzias et al., 2012; Iakoucheva et al., 2004; Marchini et al., 2011) and in prokaryotes (Singh, 2015). Ser/Thr phosphorylations within disordered stretches play crucial roles in the interactions between the proteins and their ligands. It can stabilize the tertiary structural organization of the disordered region in order to enhance its binding to the protein’s cognate ligand {Gsponer, 2008 #58}. On the other hand, once bound, it can further stabilize the bound state of the protein-ligand complex {Nishi, 2011 #59}. In addition to phosphorylation, O-linked glycosylations have also been predicted to inhabit the disordered regions of proteins (Nishikawa et al., 2010). It is reported that the evolutionary selection favored the resistance to proteolysis by coevolution of protein sequence with the site of O-glycosylation (Prates et al., 2018).

There is an emerging body of literature linking the contribution of N-linked glycosylation to the structure, tertiary fold and stabilization of proteins. In fact, intermediate steps in the biosynthesis of N-linked glycans act as checkpoints to ensure only correctly folded proteins traffic along the ER-Golgi axis (Ferris et al., 2014). Thermodynamic and molecular dynamics studies show that N-linked an O-linked glycans enhance the stability of the tertiary structure of proteins { Tams, 1998 #12; Shental-Bechor, 2008 #21; Prates, 2018 #60}. Keeping the above observations in mind, we asked whether the sites of N-glycosylations showed any bias in their locations vis-a-vis the (dis)ordered regions of N-linked glycoproteins. Investigating a dataset of 500 N-linked human glycoproteins, wherein we mapped predicted ordered and disordered regions, we found N-glycosites to be enriched predominantly within ordered regions and at some distance from the disordered stretches. We give both biochemical (proximate) and evolutionary (distal) reasons for this enrichment and argue that this mutual exclusion maintains the balance between the evolution of protein structure and function.

## Materials and Methods

### Sequence retrieval

Proteins were selected from Uniprot by sequentially applying the following criteria: 1. Encoded by the human genome and 2. Having at least one experimentally elucidated N-linked glycosylation. Subsequently the disorder prediction was performed using Genesilico Metadisorder (the best predictor of disorder based on CASP8 and CASP9) which is a metapredictor based on the following predictors of disorder: POODLE-I, IPDA, IUPRED-I, DISPRO, POODLE-S, IUPRED-S, SPRITZ-I, PRDOS, RONN, DISOPRED2, DISEMBL AND SPRITZ-S {Kozlowski, 2012 #62}. The sequences with annotated N-glycosites and predicted disordered regions is given in Supplementary File 1. Amino acid enrichment was mapped using ProtParam tool from Expasy (See Supplementary File 2) {Gasteiger, 2003 #63}.

### Gene Ontology (GO) analysis of the proteins

For prediction of gene ontology terms, sequences of the selected proteins were scanned with the GOanna server {McCarthy, 2007 #64}. GOanna performs a BLAST search against protein sequences that have a GO number.

### Phylogenetic analysis

The protein sequences were aligned using ClustalW and a phylogenetic tree was constructed using 1000 bootstrap value using MEGA5 (Tamura and Nei, 1993). Maximum likelihood was used to construct a phylogenetic tree of the proteins as it is used for the analysis of sequences of diverse origins, while maximum parsimony method was used to construct the phylogenetic tree of the proteins which have similar functions with a comparatively higher sequence homology than a former set of sequences (Tamura et al., 2011). The phylogenetic tree was viewed using the program FigTree (Rambaut, 2007). For estimation of the conservation of disorder sites and N-linked glycosylation neighborhoods, the orthologs of each protein from three species *Pan troglodytes, Mus musculus and Gallus gallus* were obtained from Ensembl and aligned with human proteins using ClustalΩ (Supplementary File 3). Shannon information entropy was measured using Protein Variability Server (http://molbiol.edu.ru/eng/index.html) (Supplementary File 4). Conservation score was calculated as follows: we assigned a score of 0 for positions where there was no conservation of residues, 0.5 where there was conservation of weakly similar residues, 1 where there was conservation of strongly similar residues and 2 where there was complete conservation of residues across the orthologs. Mean conservation of residue stretch was calculated by averaging the conservation score for the length of the residue stretch (Supplementary File 4).

### Amino acid residue frequency analysis

The frequency of amino acid residues among whole protein, disordered region and residues present in the neighborhood of N-glycosylation sites was analysed using Biostrings (Biostrings: String objects representing biological sequences, and matching algorithms, R package version 3.4.4) (Pages et al., 2009).

### Protein Functional Site prediction

In order to compute the probability that the residue combinations that are part of disordered patches or N-linked glycosylation neighborhoods and are likely to be part of, or constitute functional domains, the protein sequences were input into the Universal Evolutionary Trace server (http://lichtargelab.org/software/uet) {Lichtarge, 1996 #67}. The mean of total residue ranks was computed and compared between residues from the disordered versus N-glycosylation flanking stretches (Supplementary File 4).

### Statistics

We used both descriptive statistics and significance tests to ascertain the differences in residue characteristics between N-glycosite neighborhoods and disordered stretches. For each protein, the mean value of a given characteristic was measured across a residue stretch and when comparing across protein sets, the mean, median and standard deviation of the mean values was calculated. In order to measure significance, non-parametric Mann Whitney test was used, and when comparing across three columns, the non-parametric Kruskal-Wallis test was performed. Significance was measured through P value computed using Graphpad Prism software.

### Structure Unfolding study

In order to assay the effect of glycosylations on folding, PDB IDs of two proteins CTLA-4 (3OSK) {Yu, 2011 #71} and human pancreatic ribonuclease (2K11) {Kover, 2008 #70}were selected. In both cases, PDB files were downloaded from RCSB {Berman, 2000 #69} and GLYCAM {Kirschner, 2008 #72} was used to conjugate complex sialylated N-glycan on two distinct Asn in their sequence. Subsequently, the glycosylated and unglycosylated structures were uploaded on the MDWEB server {Hospital, 2012 #73}, and coarse-grained Brownian dynamical simulations were performed keeping criteria of Force Constant being 40 Kcal/mol* □^2^ and the distance between alpha carbon atoms of 3.8 □. The pictorial representations of the structures were obtained with PyMol.

## Results

### N-linked glycosylation sites are predominantly enriched in ordered residue stretches of N-linked glycoproteins

We used UniProt to collate a list of all human glycoproteins with at least one experimentally determined N-glycosite. The first 500 N-linked glycoproteins were processed through in Silico Metadisorder to predict the amino acids that constituted disordered sequences. This allowed us to annotate each N-linked glycoprotein for order, and N-glycosite location. We observed that 84% of experimentally determined N-linked glycosylations were predominantly located on asparagines (Asns) that were part of ordered regions rather than within the disordered stretches (Supplementary File 1; Figure 1). Most prediction and metaprediction algorithms tend to ‘overpredict’ disorder, i.e., the possibility of false positives is higher: therefore, the enrichment of N-glycan-conjugated Asns within ordered stretches could be even higher than our observation.

**Figure 1:**
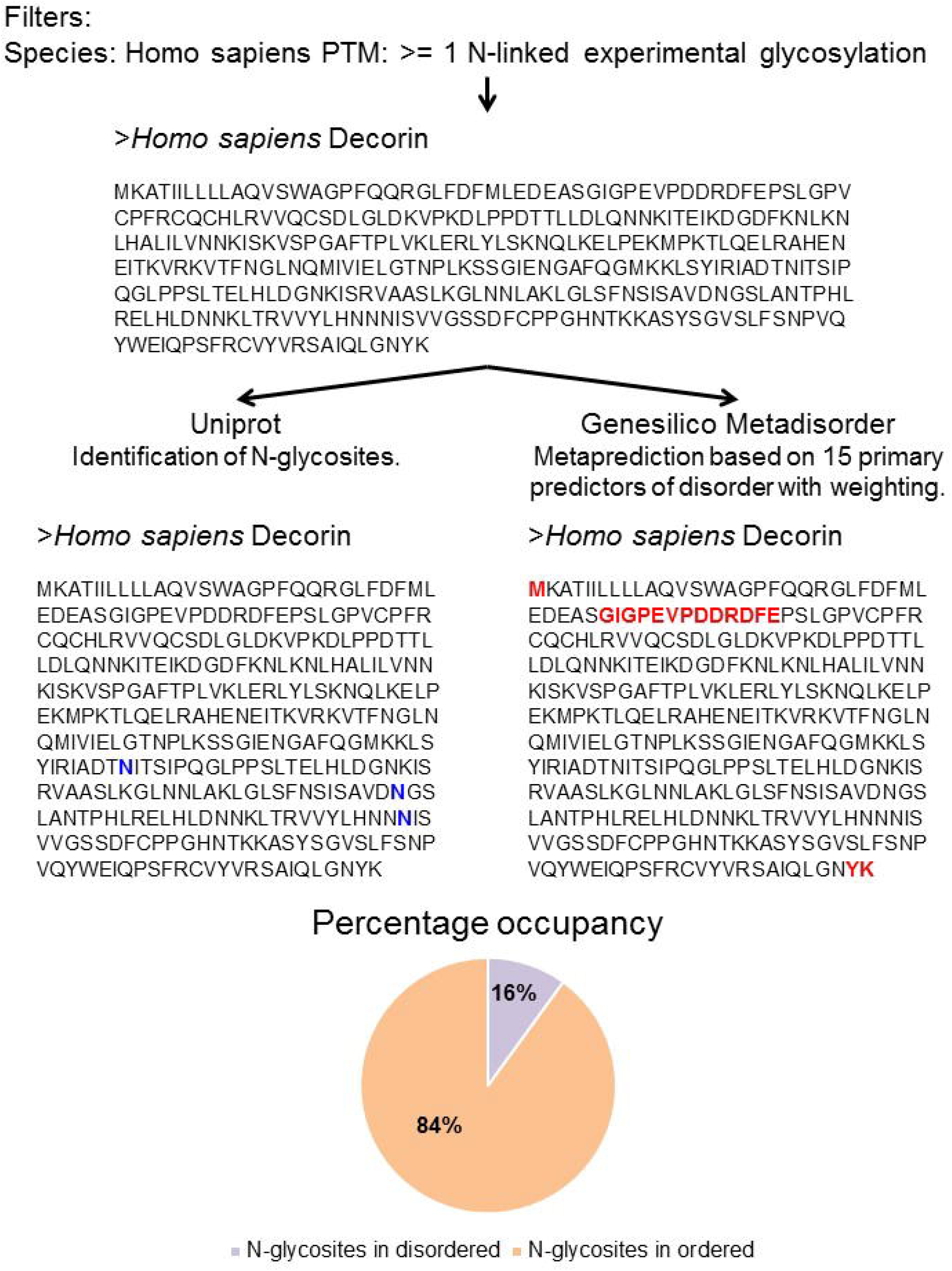
Workflow for the determination of the exclusion between N-glycosites and predicted disordered stretches. A list of proteins was compiled from Uniprot based on the criteria that they were encoded by the human genome and at the same time had atleast one N-linked glycosylation established through experimental elucidation. Subsequently, one of or more N-glycosites were annotated in its amino acid sequence. The sequence was also submitted to Genesilico Metadisorder in order to annotate the predicted disordered residue stretches. The probability of exclusion/inclusion of N-glycosites within disordered regions is shown as a pie chart (bottom). The annotaton of the N-glycosites and predicted disordered stretches is given in Supplementary File 1

In order to test the hypothesis that the exclusion of N-glycosites from disordered stretches could be cognate to a specific protein family, we analyzed with our protein set for their ontologies and localizations. We observed that within our set, glycoproteins perform a diverse range of predicted molecular and cellular functions and localized equivocally within cellular compartments (Figure S1a and b). In order to probe the ‘relatedness’ on a more stringent level (through both homology and/or evolutionary convergence, we resorted to a phylogenetic approach). The proteins were categorized on the basis of their phylogenetic proximity using both maximum likelihood and parsimony into the following categories: proteins of circulatory and skeleton system, cytoskeleton, proteins involved in the occurrence/defense of disease, nucleic acid metabolism, excretory system, hormonal system, growth factor proteins, metabolism, metalloprotein, nervous system, protein processing, reproductive system, secretory system, cellular signaling, transporter proteins, immune systems [interleukins, cytokines, tumor necrosis factor, MHC I & II, complement factor, T-cell related proteins, CD proteins, HLA class-I & II], cell adhesion molecules (selectin and integrins), polio virus receptor and antigen processing proteins (Figure S3a-h). The analysis showed that N-linked glycoproteins with disordered stretches represented a very rich and diverse set of proteins that not only clustered into several phylogenetic trees but formed nested clusters within them. This suggests that the explanation for why N-glycosites reside predominantly outside disordered regions cannot be found in the commonality of the proteins’ specific function or localization. The residue signature for N-linked glycosylation has been established to be Asn, followed by any amino acid, followed in turn by a Serine (Ser) or Threonine (Thr) (i.e., NXS/T) { Mellquist, 1998 #61}. We therefore asked whether the differential enrichment for N-glycosites could simply be reconciled by an enrichment of Asn, Ser and/or Thr within ordered stretches of proteins due to their inherent biochemical difference from disordered stretches.

### A context-specific enrichment of amino acid residues establishes the location of N-glycosites within ordered regions of N-glycoproteins

There are several descriptors of protein disorder, and one of the dominantly used descriptors is that of a consecutive stretch of polar and/or charged amino acids (Uversky et al., 2000). Since Asn, Ser and Thr are polar in nature, we estimated the abundance of these three amino acids within ordered and disordered segments of the protein dataset. The percentage of Asn was found to be not significantly different between disordered (3.7%) and ordered (4.1%) portions of N-glycoproteins. In fact, Ser and Thr were found to be significantly enriched within disordered (15.3%) regions, relative to ordered stretches (12.7%) (Table in Figure 2; Supplementary File 2). This discounted the hypothesis that the relative enrichment in N-linked glycosylation in ordered regions could be the result of a specific enrichment of Asn/Ser and/or Thr.

**Figure 2:**
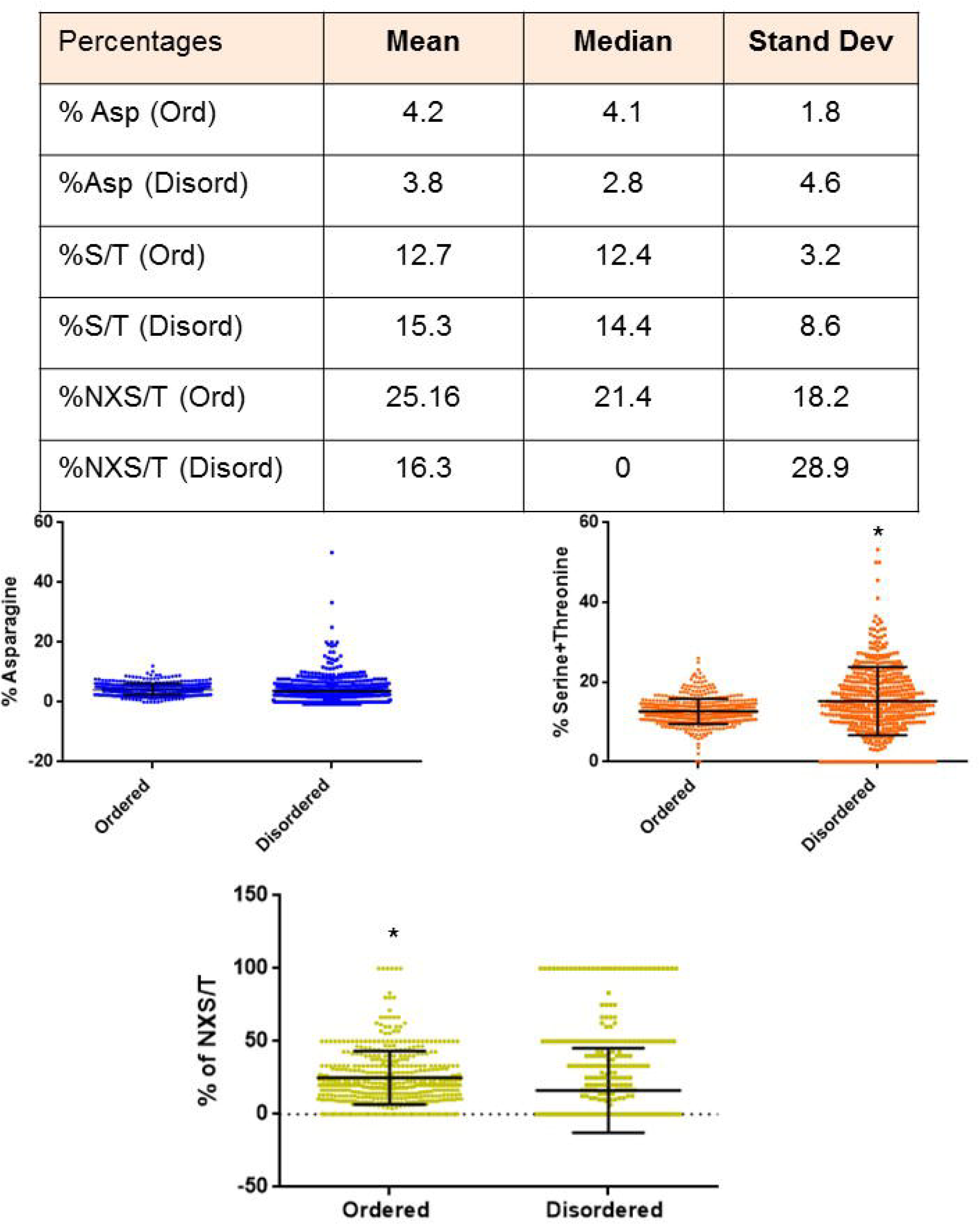
The N-linked glycosylation sequon but not the indovidual sequon residues show enrichment in the ordered regions. (Top) Table showing the mean, median and standard deviation in the percentage of Asparagines (Asn), Serines (Ser) and Threonines (Thr) and the N-linked glycosylation sequon (NXS/T) within the ordered and disordered regions of 500 human N-linked glycoproteins. (Bottom) The percentage of Asn, Ser and Thr and NXST for each protein is provided in Supplementary File 2. Graphical representation of the percentage of Asn, Ser and Thr, and NXS/T within the N-glycoprotein subset with each point representing the percentage for individual proteins. For all graphs, error bars represent SD. Statistical significance is given by P value measured using nonparametric Mann Whitney test.

We then asked what would be the probability that the amino acids, two residues downstream of any given Asn are a Ser/Thr. We observed a statistically significant divergence in the probability: 16% of Asn in the disordered regions had their second-next residue as a Ser/Thr whereas 25% of Asn in the ordered regions were followed similarly by a Ser/Thr (Figure 2). Therefore, despite an unbiased distribution of Asn between ordered and disordered regions, and a relative enrichment of Ser and Thr within the disordered region, the probability of constitution of the minimal signature required for conjugation with an N-linked glycan that consists of Asn, Ser and Thr is higher in ordered regions. We then asked if the specific amino acids that surround a given N-glycosite are conserved to a greater extent than the disordered stretches within the same N-glycoprotein regardless of its identity.

### N-glycosylation neighborhoods show evolutionary conservation within the tetrapod clade

We aligned the human N-linked glycoprotein sequences with their orthologs from chimpanzee (*Pan troglodytes*), mouse (*Mus musculus*) and chicken (*Gallus gallus*) in order to assay a possible divergence in the extent of conservation between residues surrounding the N-linked glycosylation sites (in the ordered regions) and residues in the disordered regions (Supplementary File 3). We defined N-glycosite neighborhoods as five amino acids before and after the Asn to which the N-glycan is conjugated (including the flanked NXS/T sequon). The quantification of conservation was made in two ways: first, we measured the normalized Shannon entropy of the N-glycosite neighborhoods and disordered regions (Figure 3). Shannon entropy is a measure of the information content within a sequence. Its measure across a given residue position within multiple aligned peptide sequences provides insight into how a residue is conserved in that position across species. Second, we assigned values for complete, moderate and weak conservation and averaged the individual residue score, given that disordered regions were variable in the length of their residue sequences (Figure 3). We also quantified the conservation of the minimal N-glycosite sequon (NXS/T) separately. Disordered regions show higher entropy than N-glycosite neighborhoods. The extent of residue identity conservation between the minimal signatures (i.e., NXS/T) and N-glycosylation neighborhoods was not significantly different. On the other hand, conservation of disordered residues was significantly lower than that for both the NXS/T sites as well as for N-linked glycosylation neighborhoods. Previous studies have shown that the association between residue conservation and residue disorder is context-specific {Chen, 2006 #75}{ Nishikawa, 2010 #17}. Our observation leads us to speculate that constraints in protein evolution could have operated to separate the highly conserved glycosylation sites from the relatively sparsely conserved disordered regions within glycoprotein populations. Towards this, we sought to ask whether the amino acids residues within N-glycosite neighborhoods and disordered regions are distinct in their identities. We observed that Lys showed higher abundance and Ile and Ala showed lower abundances within N-glycosite neighborhood in comparison with the whole protein sequence amino acid distribution (Figure 4 and Figure S3). Within disordered stretches, Asp showed higher abundance and Ala, Val, Ile and Arg showed lower abundance with respect to whole protein residue distribution. When compared with disordered stretches, N-glycosite neighborhoods showed higher enrichment of Val and Arg and lower enrichment of His and Phe. However, several residues showed similar enrichment between N-glycosite neighborhoods and disordered stretches such as Leu, Pro, Thr, Gly and Asp. This suggests that the residue composition of N-glycosite neighborhoods has overlap with disordered regions with key residues showing differential enrichment. We then asked whether the conservation of N-glycosite neighborhoods implicates its location with functional domains of proteins.

**Figure 3:**
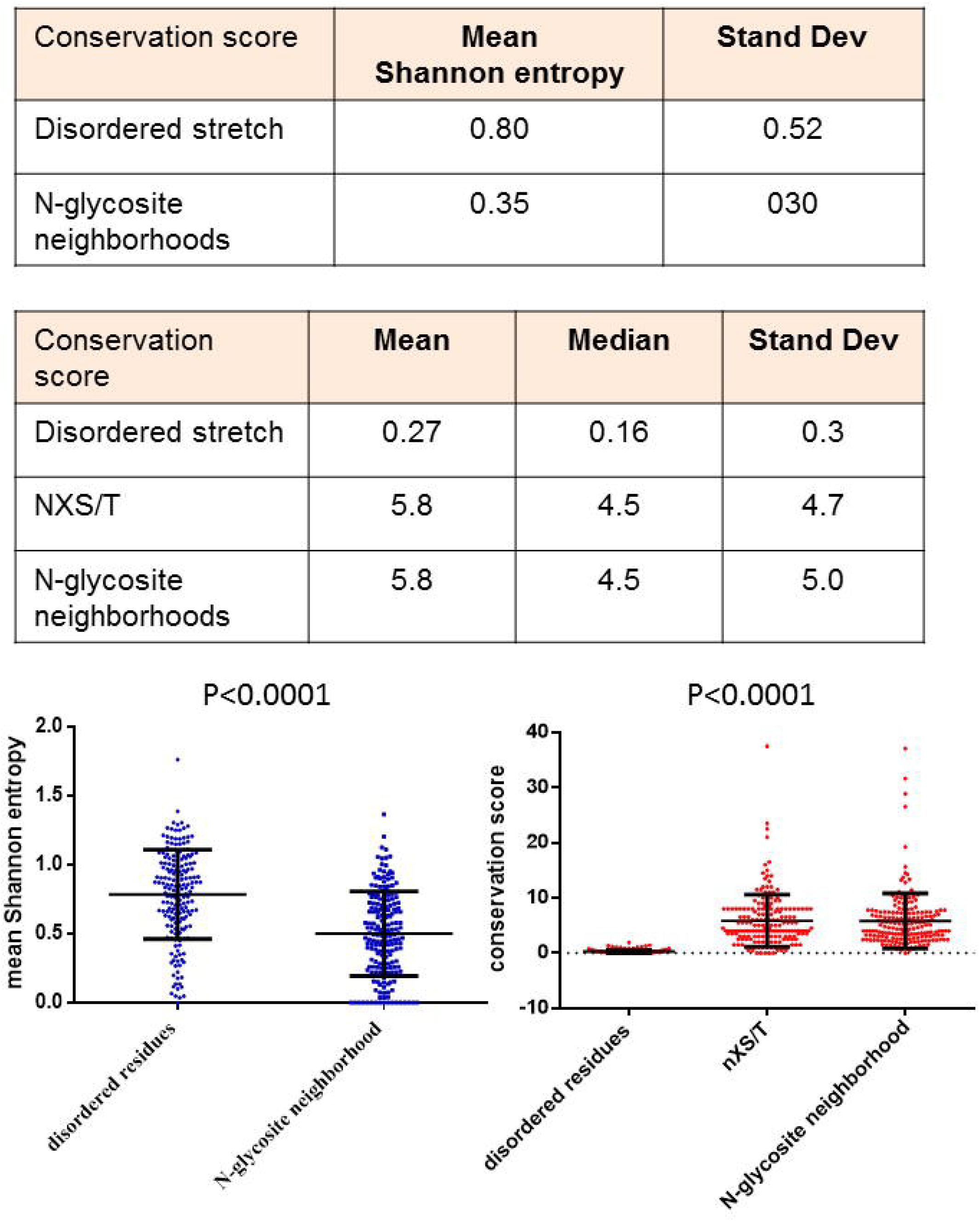
The N-linked glycosylation neighborhoods show greater phylogenetic conservation across amniote genomes than disordered residue stretches. (Top) Table showing mean Shannon information entropy of disordered residue stretches and N-glycosite neighborhoods (for definition see Main Text) and the standard deviation across N-linked glycoproteins. (Middle) Table showing mean and median conservation scores (see Methods for details on the estimation of the conservation score) of disordered stretches, NXS/T sequons and N-glycosite neighborhoods and the standard deviation within N-linked glycoproteins. (Bottom) Graphical representation of the conservation score of disordered stretches, NXS/T sequons and N-glycosite neighborhoods with each point representing the score of individual proteins. Multiple sequence alignements for each protein for which conservation was analyzed is provided in Supplementary 3. The mean Shannon entropy and computed conservation score for each protein is provided in Supplementary File 4. Error bars represent SD. Statistical significance is given by P value measured using nonparametric Mann Whitney test for two columns and Kruskal-Wallis test for three columns.

**Figure 4:**
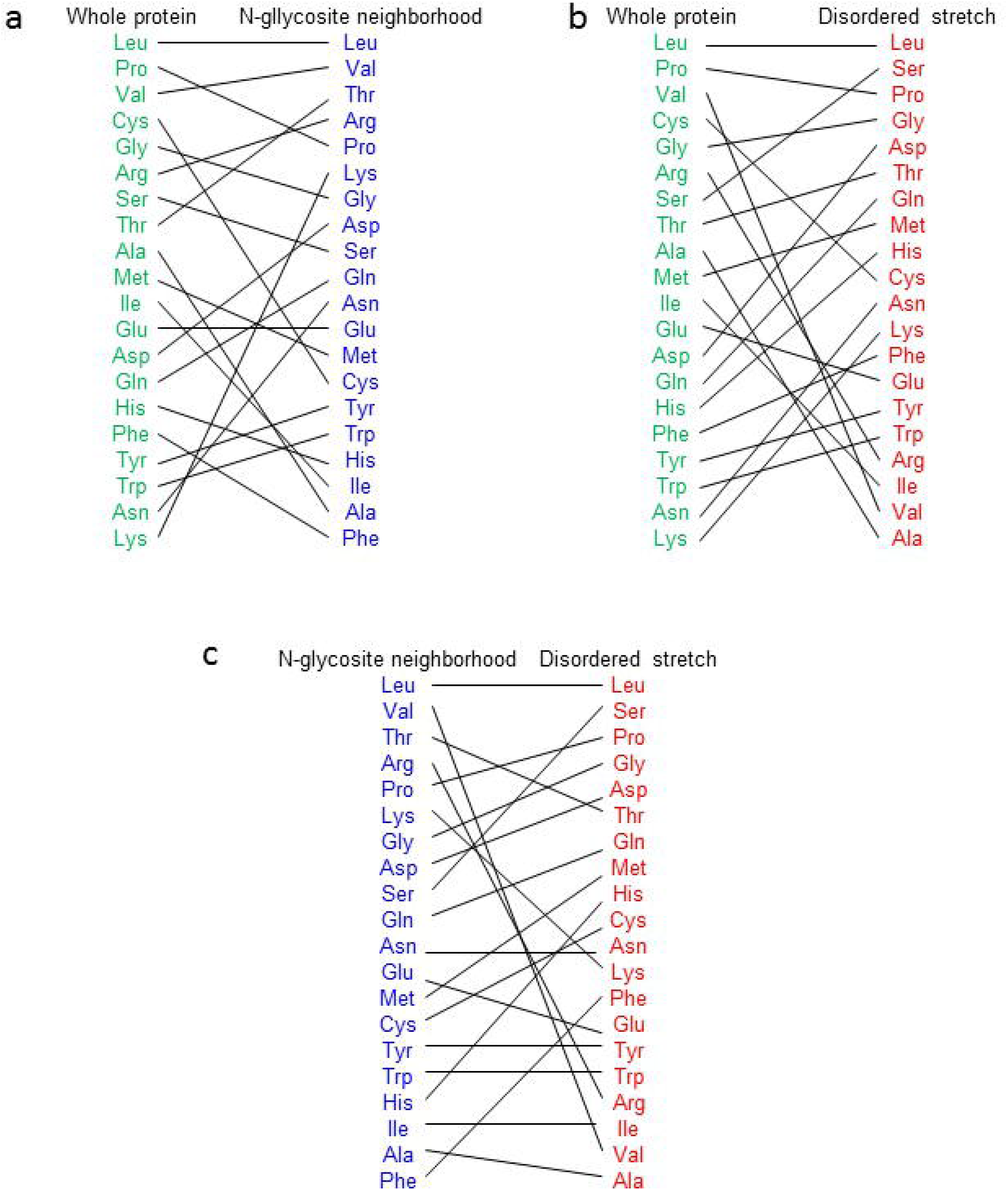
N-glycosites and disordered stretches show partially-distinct and -overlapping sets of amino acid residues. (a) Comparison of the enrichment of each amino acid within the complete protein sequence (green) and N-glycosite neighborhood (blue), arranged from high (top) to low (bottom). (b) Comparison of the enrichment of each amino acid within the complete protein sequence (green) and disordered stretches (red), arranged from high (top) to low (bottom). (c) Comparison of the enrichment of each amino acid within the N-glycosite neighborhood (blue) and disordered stretches (red), arranged from high (top) to low (bottom). The percentage of residue abundance is given in Figure S3.

### Disordered stretches show greater potential than N-glycosite neighborhoods to be part of functional protein domains

We used evolutionary trace, an algorithm that computes the functional importance of each amino acid within a protein based on variation within closely- and distantly related species {Lichtarge, 1996 #67}. The functional importance is therefore not centered on conservation but on the fact that variation in distant species most likely altered protein function and rendered the amino acid functionally important. Such important amino acids potentially cluster together allowing prediction of putative functional domains for proteins that have not been crystallized. To our surprise, the disordered stretches of N-linked glycoproteins showed greater clustering of amino acids with significantly high functional importance relative to their N-glycosite neighborhoods (Figure 5). This suggests that N-glycosite neighborhoods while evolutionarily conserved, show a relatively lower tendency to be part of functional protein domains.

**Figure 5:**
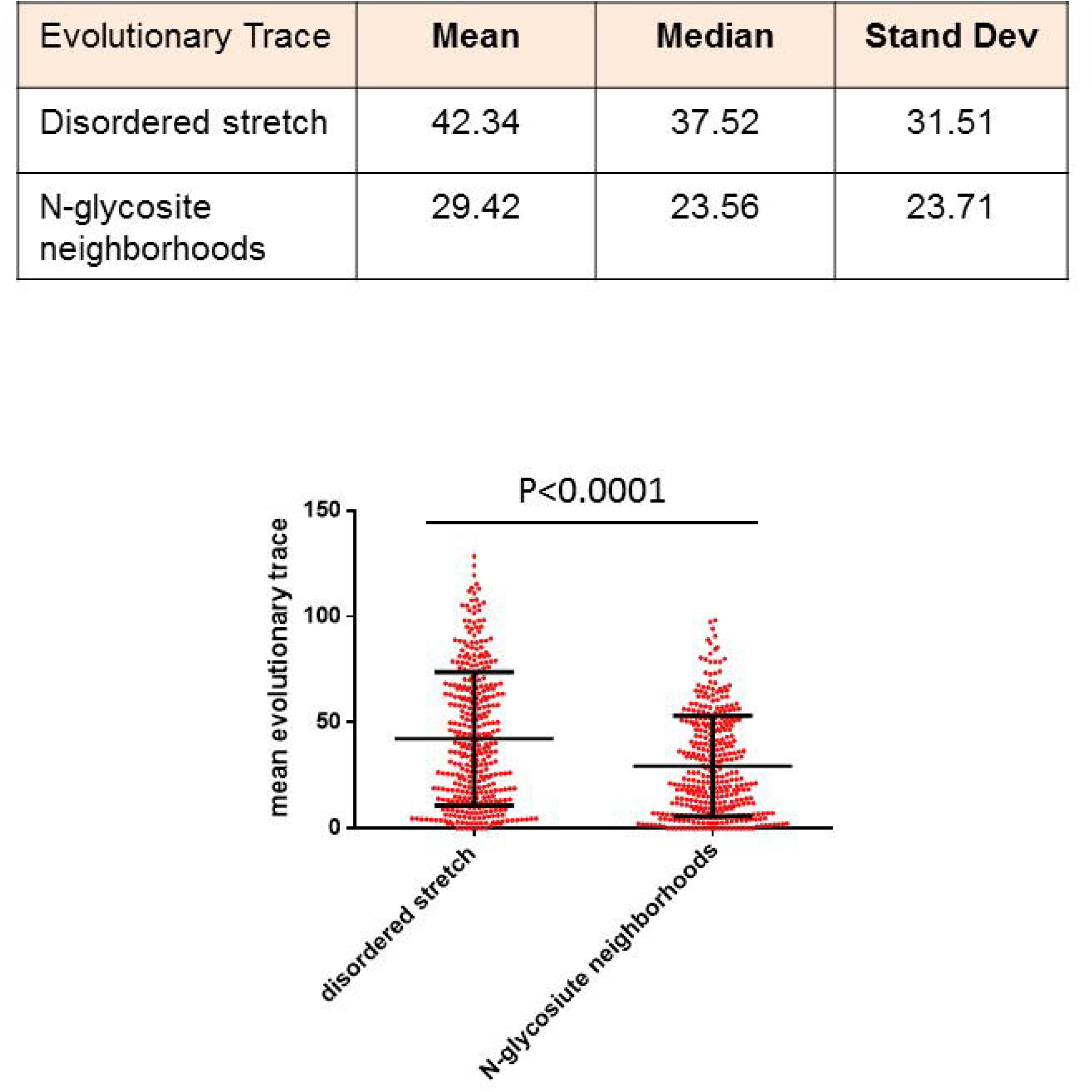
The N-linked glycosylation neighborhoods show poorer ability to be part of functional domains in comparison with diordered residue stretches. (Top) Table showing mean and median Evolutionary Trace of disordered residue stretches and N-glycosite neighborhoods (for definition see Main Text) and the standard deviation across N-linked glycoproteins. (Bottom) Graphical representation of the evolutionary traces of disordered stretches and N-glycosite neighborhoods with each point representing the score of individual proteins. The mean evolutionary trace for each protein is provided in Supplementary File 4. Error bars represent SD. Statistical significance is given by P value measured using nonparametric Mann Whitney test.

## Discussion

An emerging literature proposes the tradeoff between the evolution of protein sequence, structure and function (thermodynamic stability and folding) (Gosavi, 2013; Schreiber et al., 1994; Shoichet et al., 1995). Functional residues that are part of binding and interacting interfaces potentially hinder folding and affect the stability of the tertiary structure. The intuitive explanation for this tradeoff is that the exposed residues that can be subject to evolutionary bias in order to increase the folding of the protein cannot be the relevant residues for the functioning of the proteins or to be precise, a given protein domain. Understanding the parsing of structure- and function-contributing residues represents a fascinating problem of molecular evolution or conservation pattern followed by amino acid residues.

The protein sequences used in this study were found to be involved in different biological and molecular functions (Figure S1). In addition, the phylogenetic analysis does not provide any direct evidence about the pattern of distribution of disordered regions in the proteins (Figure S2). However, it was observed that proteins which shared some common function or related to the same cellular processes, often were clustered in one group. This clustering of proteins provides a basic idea about the evolutionary relatedness among them. Therefore, these may help to choose representatives of clustered protein sets for further studies. Disordered regions are known to contain variable amino acid sequences, and we observe that they show relatively low phylogenetic conservation even among amniote genomes. Despite this, some amino acid residues which have longer side chains than others e.g. serine, threonine, aspartate, glutamine, and leucine are present repeatedly in the disordered region. The long chains of these amino acids may have some significant function in the disordered region which could be related either to bury or expose the amino acids according to their position. However, amino acid residues which have small side chain e.g. glycine is also abundant in whole protein and disordered regions. This is attributed to its ability to be present in the maximum conformational states allowed by Ramachandran plot and its small size allows it to fit into the β-turns constituting the structure of the protein. Proline also follow the same pattern as glycine, however, its abundance may be due to the presence of pyrrolidine ring which contain imino group present in its side chain, which allows it to introduce kinks or turns in the protein structure. It is well known that high proline content is generally found in disordered proteins {Prates, 2018 #60}. Our results from this study lead us to propose that a spatial separation between sites of N-glycosites and protein disordered region sites potentially acts as a constraint to distinguish residue stretches that contribute to protein structure and function. This proposition is based on the mounting theoretical (Lee et al., 2015; Shental-Bechor and Levy, 2008) and experimental (Banks, 2011) evidence that glycosylations impact the folding and structure of glycoproteins (although the relationship is not always positively regulatory) (Gavrilov et al., 2015; Shental-Bechor and Levy, 2011)). The triose core of N-glycoproteins enhances both the kinetics and stability of tertiary glycoprotein folds (Hanson et al., 2009). In fact, N-glycosylations seems to destabilize the unfolded state more than the folded state of the protein (Shental-Bechor and Levy, 2008). It follows, therefore, from these studies that where the protein possesses a particular region which needs to be disordered to act as an interacting and binding interface, contributing to the functionality of the protein, the stability-providing sites of N-glycosylations should remain spatially excluded from such disordered regions. To test our hypothesis, we have examined the Brownian dynamics of two glycoproteins cytotoxic T-lymphocyte antigen 4 (CTLA-4) (RCSB ID: 3OSK) and human pancreatic ribonuclease (RCSB ID: 2k11) with or without conjugation with a complex sialylated glycan geometry. In confirmation with our hypothesis, both unconjugated proteins unfolded much faster than glycoconjugated counterparts.

The N-linked glycosylation of proteins also tends to act as an important quality control check point for the ability of the protein to fold itself correctly (Xu and Ng, 2015). Unfolded proteins are N-glycosylated within the ER lumen and bear 3 Glc residues at the terminal end of their A-branch. N-linked glycans being bulky hydrophilic polymers can increase the thermodynamic stability of proteins, enhancing their folding. This is temporally coincident with the cleavage of Glc by glucosidases. The de-glucosylated N-glycoproteins, if not folded properly, are bound by an enzyme UGGT which binds to them and re-glucosylates restarting the process of deglucosylation-associated folding. Failure of folding results in the proteolytic degradation of unfolded proteins whereas folded proteins escape detection by UGGT allowing the latter to be a glycosylation-based checkpoint that examines the folded state of glycoproteins (Sifers, 2004). The presence of an N-linked glycosite within a disordered stretch of a glycoprotein would potentially foreclose the possibility of its evasion of the UGGT checkpoint resulting in the degradation of such proteins. We believe this could be an important reason for the separation and evolutionary stabilization of N linked glycosites from disordered regions of the N-glycoprotein.

It is pertinent to ask whether the biochemical properties of the amino acids that constitute the disordered stretches are dissimilar to those that surround the N-glycosylation sites. One of the widely studied biochemical defining features of disordered proteins is the polar, hydrophilic amino acids that comprise them. Uversky and coworkers have even come up with a charge-hydropathy plots wherein a separatrix can be drawn between ordered and disordered protein clusters (Oldfield et al., 2005). We were not able to seek a clear separation between N-glycosylation neighborhoods and disordered regions using such plots. Our results showed that while Asparagine, Serine or Threonine were not significantly enriched within ordered regions of N-linked glycoproteins, the NXS/T signature was enriched within such regions. The occurrence of glycosylation-permissive signatures could result in the disruption of the function by affecting the ‘unstructuredness’ within disordered stretches and were potentially weeded out through selection.

Our results are specific to N-linked glycoproteins which are part of the extracellular milieu. Many of them being transmembrane- and secreted diffusible-proteins, function across multicellular tissue spatial scales and contribute to the developmental mechanisms by mediating discrete biophysical and biochemical functions (Newman and Bhat, 2009). The determinacy of their developmental roles is a function of their stability and N-glycosylation may have played a very important role in this. Our study qualifies this by proposing that the localization of N-linked glycosylations allowed these proteins to also have unstructured interfaces in order that they interact with the various components in the extracellular milieu without compromising the structured fold of the proteins. Based on the present phylogenetic analysis of the proteins, select proteins can be taken as representatives and the role of disordered regions can be studied in detail by molecular dynamics-based approaches. Further studies will help better understand the role of disordered regions in the functionality of proteins at the molecular level.

---

**Figure S1:** N-linked Glycosylated sites selected proteins with GO (Biological, cellular and molecular function) terms.

**Figure S2: (a) Phylogenetic tree of proteins belonging to circulatory syatem**: Proteins clustered into groups depending on their cellular function: Proteins shown in blue colour belong to coagulation factors, magenta colour encode protein involved in gas transport, protein showed in red colour are related to regulation of blood cells. **(b) Phylogenetic tree of cytosekeletal proteins:** Cytoskeletal proteins formed different groups according to their functions (blue-Collagenase), maintaining cell shape (Red-Microfibrillar and cell adhesion protein), muscle related protein (green-sarcoglycan) and protein of extracellular matrix (Black). **(c) Phylogenetic relationship among proteins of nucleic acid metabolism** Proteins of nucleic acid metabolism clustered according to their function: proteins which act on the outer side of DNA encoded in black, proteins which cleave nucleic acid encoded in red. **(d) Proteins containing disordered region belong to different functional classes of the immune system:** It was observed that the proteins get clustered into different groups depending on their function or their interaction with the antigen/ antigen processing step. The proteins showed in red colour code for the proteins related to Immunoglobulins, antigen-related cell adhesion molecule, selectin protein, purple colour encode HLA class I histocompatibility antigen for alpha chain and different proteins which involve glycoprotein interactions like CTLA, Neutrophil gelatinase-associated lipocalin (NGAL), cyan blue colour encode HLA class I histocompatibility antigen a or C class for alpha chain, wheat brown colour encode for Antigen-presenting glycoprotein CD proteins and some HLA class I histocompatibility antigen DR beta chain, black colour encode proteins related to CD like CD63, CD244 Natural killer cell receptor, CD80 T-lymphocyte activation antigen, CSF2R Granulocyte-macrophage colony-stimulating factor receptor which exhibit different functions, dark blue colour encode for different proteins ranging from interferon to proteins involved in naive B-cell development, complement factor related protein, interleukins and Lysosome-associated membrane glycoprotein, green colour encode Immunoglobulin heavy constant gamma protein while IL17 and IL2 also showed similarity to these proteins and some miscellaneous proteins formed a separate showed in black colour are Ficolin-3, Killer cell immunoglobulin-like receptor, Leukocyte immunoglobulin-like receptor, Mucosal addressin cell adhesion molecule, Epithelial cell adhesion molecule Tubulointerstitial nephritis antigen-like and Inhibitor of nuclear factor kappa-B kinase-interacting protein. Most of the proteins taken for the study belong to the immune system and formed groups according to their function. **(e) Proteins involved in the metabolic processes segregated according to their role in different metabolic activities:** The proteins involved in the removal of phosphate from glucose formed one group shown in blue colour. ER protein represented in cyan blue colour and proteins involved in galactose processing were shown in magenta colour. Serum paraoxonase/arylesterase proteins formed a cluster shown in yellow colour. Antitrypsin and antichymotrysin formed cluster presented in red colour. Glucoside xylosyltransferase and Lysosomal acid phosphatase showed similarity and presented in green colour. Protein related protein digestion, fat metabolism and gastric digestion regulation formed a cluster shown in purple colour. Apolipoprotein C-IV, Glutaminyl-peptide cyclotransferase and Exostosin-like 2 formed a cluster which is shown in dark blue colour. Rest of the protein represented in black colour belong to different categories like Dipeptidase, Gastric intrinsic factor, Plasma alpha-L-fucosidase, Thioredoxin domain-containing protein, Lysosomal thioesterase, Calcium homeostasis modulator protein etc. did not form a separate cluster as they have different function and there is not much similarity between their sequences. **(f) Phylogenetic tree of proteins involved in its processing:** Transmembrane emp24 domain-containing protein formed a cluster shown in purple colour. Endoplasmic reticulum-Golgi intermediate compartment protein showed in red colour. Protein disulfide-isomerase, Calumenin protein which is present in ER, Signal peptidase complex subunit, Di-N-acetylchitobiase, Translocon-associated protein and Sulfatase-modifying factor formed a cluster shown in cyan blue colour. Cation-dependent mannose-6-phosphate receptor and Nucleotide exchange factor protein belong to the different class of proteins, however, they showed similarity and presented inblue colour. Rest of the proteins shown in black did not show similarity based on their function and sequence. **(g) Protein related to reproduction clustered based on their regulatory role in the system:** Lutropin subunit β, Follistatin-related protein, Basigin, Protein LEG1, Cysteine-rich secretory protein,and Glycodelin formed one cluster shown in magenta colour. All of these proteins are either involved in the regulation of early development, growth, pubertal maturation, and reproductive processes. Indian hedgehog protein, Sex hormone-binding globulin, Alpha-2-HS-glycoprotein, Inhibin alpha chain and Prostatic acid phosphatase are involved in different functions and did not show any homology in their sequence. **(h) Proteins involved in the transport of biomolecules showed homology based on their function:** Sodium/potassium-transporting ATPase, ATP-sensitive inward rectifier potassium channel and Ileal sodium/bile acid cotransporter shared a similarity in sequence and represented in purple colour. Equilibrative nucleoside transporter are presented in blue colour. Glycine receptor subunit alpha-3 and lymphatic vessel endothelial hyaluronic acid receptor showed similarity which is shown in cyan blue colour. Alpha-1-acid glycoprotein and Apolipoprotein D formed a cluster shown in green colour. Potassium channel subfamily K formed a separate cluster shown in red colour. Rest of the proteins did not follow and show sequence similarities like Transthyretin, Translocon-associated protein Glycolipid transfer protein, Calcium-activated potassium channel, Folate receptor, Leucine-rich repeat-containing protein and Ganglioside GM2 activator.

**Figure S3: Estimation of frequency of amino acids within whole protein sequences (top), disordered stretched (middle) and N-glycosite neighborhoods (bottom)**

**Figure S4: Brownian dynamics of N-linked glycosylated proteins with disorder shows slower unfolding in presence of glycosylation.** (A) PyMol-derived structures of human cytotoxic T lymphocyte Antigen-4 (CTLA-4 RCSB ID: 3OSK) with no N-linked glycosylation (left), a complex glycan tree conjugated on Asn78 (middle) or Asn110 (right) (glycosylation added using Glycam). (B) Graph showing the variation in radius of gyration along a trajectory of Brownian dynamics for the three structures of CTLA4 described in (A). Black trace shows the temporal dynamics of gyration of unglycosylated CTLA4 whereas the red and orange traces show the temporal dynamics of gyration of CTLA4 glycosylated at Asn78 and Asn110 respectively. (C) PyMol-derived structures of human pancreatic ribonuclease1 (RNASE1 RCSB ID: 2K11) with no N-linked glycosylation (left), a complex glycan tree conjugated on Asn34 (middle) or Asn76 (right) (glycosylation added using Glycam). (B) Graph showing the variation in radius of gyration along a trajectory of Brownian dynamics for the three structures of 2K11 described in (A). Black trace shows the temporal dynamics of gyration of unglycosylated 2K11, whereas the red and orange traces show the temporal dynamics of gyration of 2K11 glycosylated at Asn34 and Asn76 respectively.

